# Yeasts have evolved divergent enzyme strategies to deconstruct and metabolize xylan

**DOI:** 10.1101/2022.12.22.521714

**Authors:** Jonas L. Ravn, Amanda Sörensen Ristinmaa, Tom Coleman, Johan Larsbrink, Cecilia Geijer

## Abstract

Together with bacteria and fungi, yeasts actively take part in the global carbon cycle. Over a hundred yeast species have been shown to grow on the major plant polysaccharide xylan, which requires an arsenal of carbohydrate active enzymes. However, which enzymatic strategies yeasts use to deconstruct xylan and what specific biological roles they play in its conversion remain unclear. In fact, genome analyses reveal that many xylan-metabolizing yeasts lack expected xylanolytic enzymes. Guided by bioinformatics, we have here selected three xylan-metabolizing ascomycetous yeasts for in-depth characterization of growth behavior and xylanolytic enzymes. The savanna soil yeast *Blastobotrys mokoenaii* displays superior growth on xylan thanks to an efficient secreted glycoside hydrolase family 11 (GH11) xylanase; solving its crystal structure revealed a high similarity to xylanases from filamentous fungi. The termite gut-associated *Scheffersomyces lignosus* in contrast grows more slowly and its xylanase activity was found to be mainly cell surface-associated. The wood-isolated *Wickerhamomyces canadensis* surprisingly could not utilize xylan as the sole carbon source without adding xylooligosaccharides, exogenous xylanases or even by co-culturing with *B. mokoenaii*, suggesting that *W. canadensis* relies on initial xylan hydrolysis by neighboring cells. Furthermore, our characterization of a novel *W. canadensis* GH5 subfamily 49 (GH5_49) xylanase represents the first demonstrated activity in this subfamily. Our collective results provide new information on the variable xylanolytic systems evolved by yeasts and their potential roles in natural carbohydrate conversion.

**Importance:** Microbes that take part in the degradation of the polysaccharide xylan, the major hemicellulose component in plant biomass, are equipped with specialized enzyme machineries to hydrolyze the polymer into monosaccharides for further metabolism. However, despite being found in virtually every habitat, little is known of how yeasts break down and metabolize xylan and what biological role they may play in its turnover in nature. Here, we have explored the enzymatic xylan deconstruction strategies of three underexplored yeasts from diverse environments: *Blastobotrys mokoenaii* from soil, *Scheffersomyces lignosus* from insect guts and *Wickerhamomyces canadensis* from trees, and show that each species has a distinct behavior regarding xylan conversion. These findings may be of high relevance for future design and development of microbial cell factories and biorefineries utilizing renewable plant biomass.

## Introduction

Yeasts inhabit every biome in nature and live in a variety of ecological niches including soil, water, air, and on plant and fruit surfaces (1). Together with filamentous fungi and bacteria, yeasts actively partake in polysaccharide decomposition in decaying plant biomass but have been much less studied. In a recent review on cellulose and xylan-degrading yeasts, we concluded that cellulose-degrading yeasts are rare whereas xylan-degrading yeasts are rather widespread in nature. In fact, more than 100 yeast species have been identified to date that displays xylanolytic capacities, but their precise xylan degradation strategies and biological role(s) in the ecosystem remain largely unexplored (2).

Microorganisms use Carbohydrate Active Enzymes (CAZymes), collected and described within the CAZy database (www.cazy.org; (3), to degrade xylan polymers into monosaccharides that can be further catabolized and used as carbon and energy source. Xylans are comprised of a backbone of β-1,4-linked D-xylose residues that may be highly *O*-acylated and substituted by α-1,2-linked (methyl)-glucuronic acid, α-1,2- or α-1,3-linked arabinosyl units and phenolic compounds (4, 5). These decorations give rise to different types of xylans typically grouped into arabinoxylan (AX), glucuronoxylan (GX), and glucuronoarabinoxylan (GAX) (6). Xylan degradation requires an arsenal of CAZymes such as carbohydrate esterases (CEs) and glycoside hydrolases (GHs) with the most common xylan backbone-cleaving *endo*-β-1,4-xylanases being found in glycoside hydrolase family 10 (GH10), GH11, and GH30 (7, 8).

In a previous study, we predicted CAZymes in 332 genome-sequenced ascomycetous yeasts (9) and identified several new xylanolytic species (10). Combined with additional sequenced xylanolytic yeasts in literature, 24 species and their respective CAZymes were bioinformatically mapped (2). Interestingly, we identified different subgroups of xylanolytic yeasts regarding their putative *endo*-β-1,4-xylanases. Only a single species encodes a GH11 xylanase, while GH10 and GH30_7 enzymes were encoded by five and three species, respectively. However, 18 out of the 24 yeasts lack enzymes from those common xylanase families, indicating that they possess novel xylanases or xylanolytic strategies (2, 10). Many encode GH5 enzymes from subfamilies without known specificities and we hypothesize that these may be missing pieces in the xylan deconstruction puzzle of yeasts (Fig. 1).

**Figure 1.**
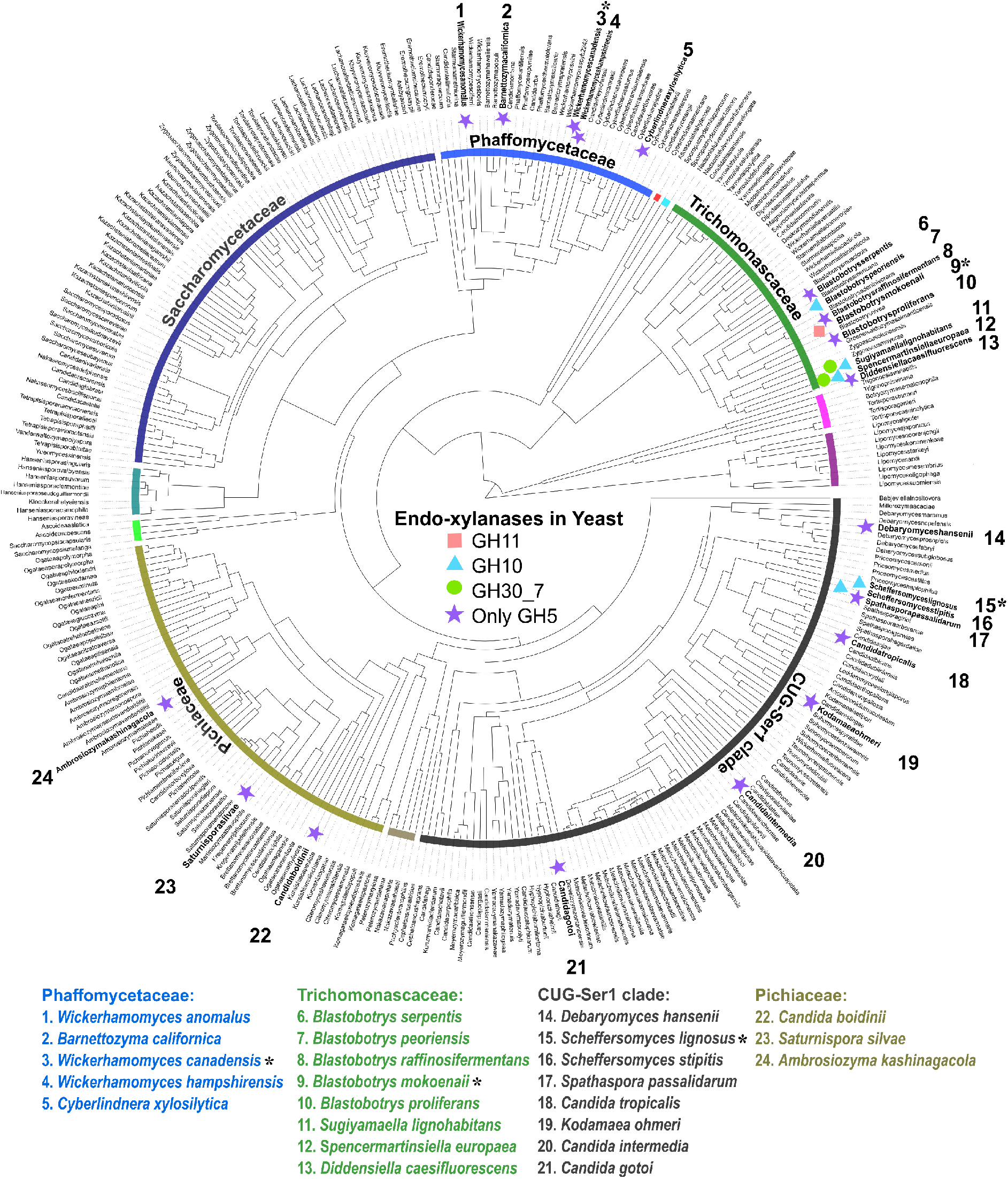
Overview of predicted endo-xylanases in known xylanolytic yeasts. (A) Putative xylanases are marked next to known xylanolytic yeasts (highlighted bold and by numbers). Asterixis (*) indicate the three yeasts selected for characterization. GH = glycoside hydrolase.

In this study, we characterized the xylanolytic strategies of three phylogenetically diverse yeasts with different CAZyme profiles (10): *Blastobotrys mokoenaii* (phylogenetically in the *Trichomonascaceae* clade; isolated from savanna soil) which encodes a GH11 and a GH30_7 xylanase, *Scheffersomyces lignosus* (CUG-Ser1 clade; wood-boring insects) which encodes a GH10 xylanase, and *Wickerhamomyces canadensis* (*Phaffomycetaceae* clade; Canadian Red Pine) which lacks any obvious *endo*-β-1,4-xylanases. We successfully applied a strategy based on a rational bioinformatic selection of enzyme targets, heterologous production, and screening of recombinant enzymes on a wide diversity of carbohydrates, which confirmed xylanase activities of *Bm*GH11 and *Sl*GH10. Moreover, we demonstrated that *W. canadensis* possesses functional GH5 subfamily 49 (GH5_49) and GH5_22 *endo*-xylanases.

## Results

### Yeast xylan growth and subcellular localization of xylanolytic activities

To characterize the xylanolytic growth behavior of *B. mokoenaii, S. lignosus*, and *W. canadensis*, we monitored their growth in minimal medium with beechwood GX or xylose as sole carbon sources. *B. mokoenaii* reached a relatively high final OD_600_ (~14) on beechwood GX with similar growth characteristics as on xylose. In contrast to the two other species with typical spherical yeast shapes (Fig. 2E-F), *B. mokoenaii* displayed pseudo-mycelia (Fig. 2D) and multilateral budding morphology forming hyphea and blastoconidia with elongated setae (Kurtzman and Robnett 2007; Mokwena et al., 2000). *S. lignosus* displayed an extended lag-phase and slower growth on beechwood GX compared to xylose, while no growth of *W. canadensis* on xylan could be detected within the timeframe of the experiment (96 h). We hypothesized that the lack of growth may be attributed to lack of sensing the carbohydrate polymer and decided to supplement the medium with 0.2 % xylooligosaccharides (XOs), which enabled *W. canadensis* to grow to an OD_600_ of ~5. As 0.2 % XOs alone did not support significant growth (OD ~1), this strongly suggests that addition of XOs successfully induced a functional xylanolytic system of this species (Fig. 2C).

**Figure 2.**
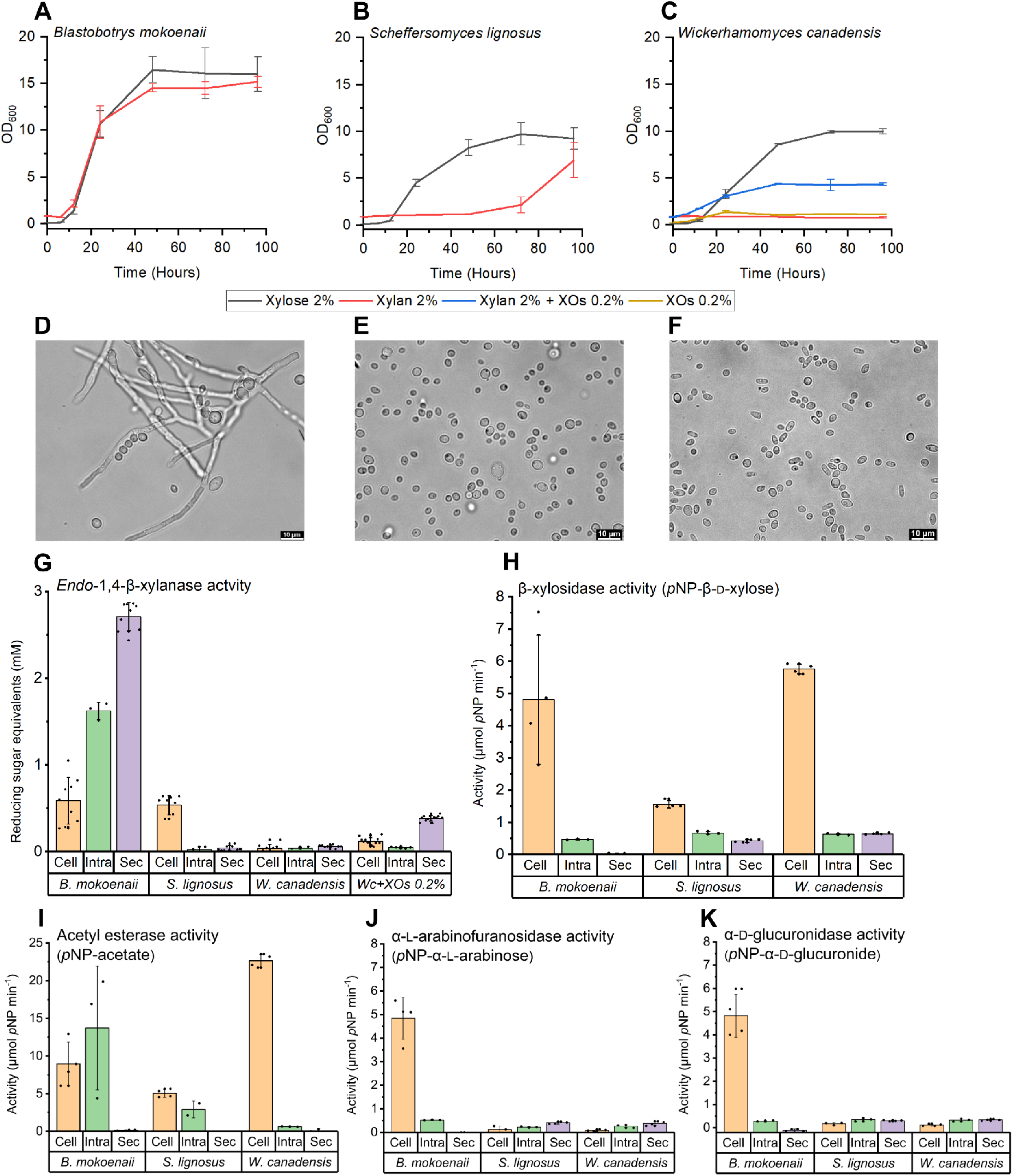
Yeast growth in beechwood glucuronoxylan and xylanolytic activity localization. (A-C) Yeasts were grown in 10 mL Delft minimal medium with 2 % beechwood glucuronoxylan or 2 % xylose as sole carbon source in biological triplicates. *Wc* = *Wickerhamomyces canadensis;* XOs = xylooligosaccharides. (D-F) Brightfield microscopy showing yeast morphology from 96 h xylan cultures. (G) Yeast subcellular xylanase activity originating from secretome (Sec), cells (Cell) and intracellular (Intra) from lysed cell fractions were compared using DNS reducing sugar assays in triplicates. (H) β-xylosidase activity was quantified using *p*-nitrophenyl-β-D-xylopyranoside, (I) acetyl esterase activity using *p*-nitrophenyl-acetate, (J) α-L-arabinofuranosidase activity using *p*-nitrophenyl-α-L-arabinofuranose, and (K) α-D-glucuronidase activity using *p*-nitrophenyl-α-D-glucuronide.

*Endo*-β-1,4-xylanases are the main enzymes responsible for xylan depolymerization, supported by β-xylosidases. The enzymes can either be secreted extracellularly, attached to the cell surface, or intracellular. To determine the subcellular localization of the enzymes in the three yeasts, the activities in the secretome, intact live yeast cells and lysed cells were determined by the dinitrosalicylic acid (DNS) assay to detect released reducing sugars resulting from xylan cleavage. β-xylosidase activities were determined using *p*-nitrophenyl-β-D-xylopyranoside. Clear extracellular xylanase activity was observed for *B. mokoenaii*, while for *S. lignosus* xylanolytic activity appeared to be mainly cell surface-associated. *W. canadensis* cells that had been incubated on xylan alone exhibited little or no activity, while cells exposed to XOs and xylan had extracellular xylanase activity (Fig. 2G). β-xylosidase activity was mainly cell surface-associated in all three yeasts (Fig. 2H), as were other accessory activities such as acetyl esterases (Fig. 2I). Only *B. mokoenaii* showed significant cell-associated α-L-arabinofuranosidase and α-D-glucuronidase activity (Fig. 2J-K), which correlates well with the predicted CAZyme profiles for the three yeasts where only *B. mokoenaii* possesses putative α-L-arabinofuranosidases (GH43, 51, *62)* and an α-D-glucuronidase (GH67). *S. lignosus* possesses another putative α-D-glucuronidase (GH115), while *W. canadensis* appears to lack these enzymatic activities altogether (10). Overall, these results conclude that all three yeasts can degrade xylan polymers and use the released sugars for growth, likely through the combined action of *endo*-β-1,4-xylanases, β-xylosidases and other accessory enzymes. However, as evident from the differences in growth profiles and the enzyme assays, the three species use different xylanolytic strategies for xylan degradation.

### Biological roles of the different yeast species

The fact that addition of XOs boosts *W. canadensis’* growth on xylan indicates that, in nature, this species relies on other microorganisms to initiate the hydrolysis of intact xylan polymers. Secreted enzymes such as the GH11 from *B. mokoenaii* produce XOs in the extracellular environment, which could also benefit neighboring cells from other species and enable cross-feeding behavior. We co-cultured *W. canadensis* and *B. mokoenaii*, which interestingly led to synergistic growth behavior, with higher final ODs (at 120 h) than in both respective monocultures (Fig. 3A). Moreover, we were able to utilize different fluorescent staining patterns (see Fig. S1 for controls) of the two species using Calcofluor white and FUN-1, to follow their individual growth throughout the co-culture (Fig. 3B-E). *B. mokoenaii* appears to supply *W. canadensis* with XOs by its secreted *endo*-xylanase, which could also be observed on agar plates, where *B. mokoenaii* created a clearing zone from xylan hydrolysis that upon reaching *W. canadensis* enabled growth (Fig. 3F-G and supplemental gifs (link) for a 21-day time lapse). These results correlate well with the *W. canadensis* growth induced by addition of XO as described above, and strongly indicate that externally supplied xylanase activity triggers *W. canadensis* growth on xylan and expression of its own xylanases.

**Figure 3.**
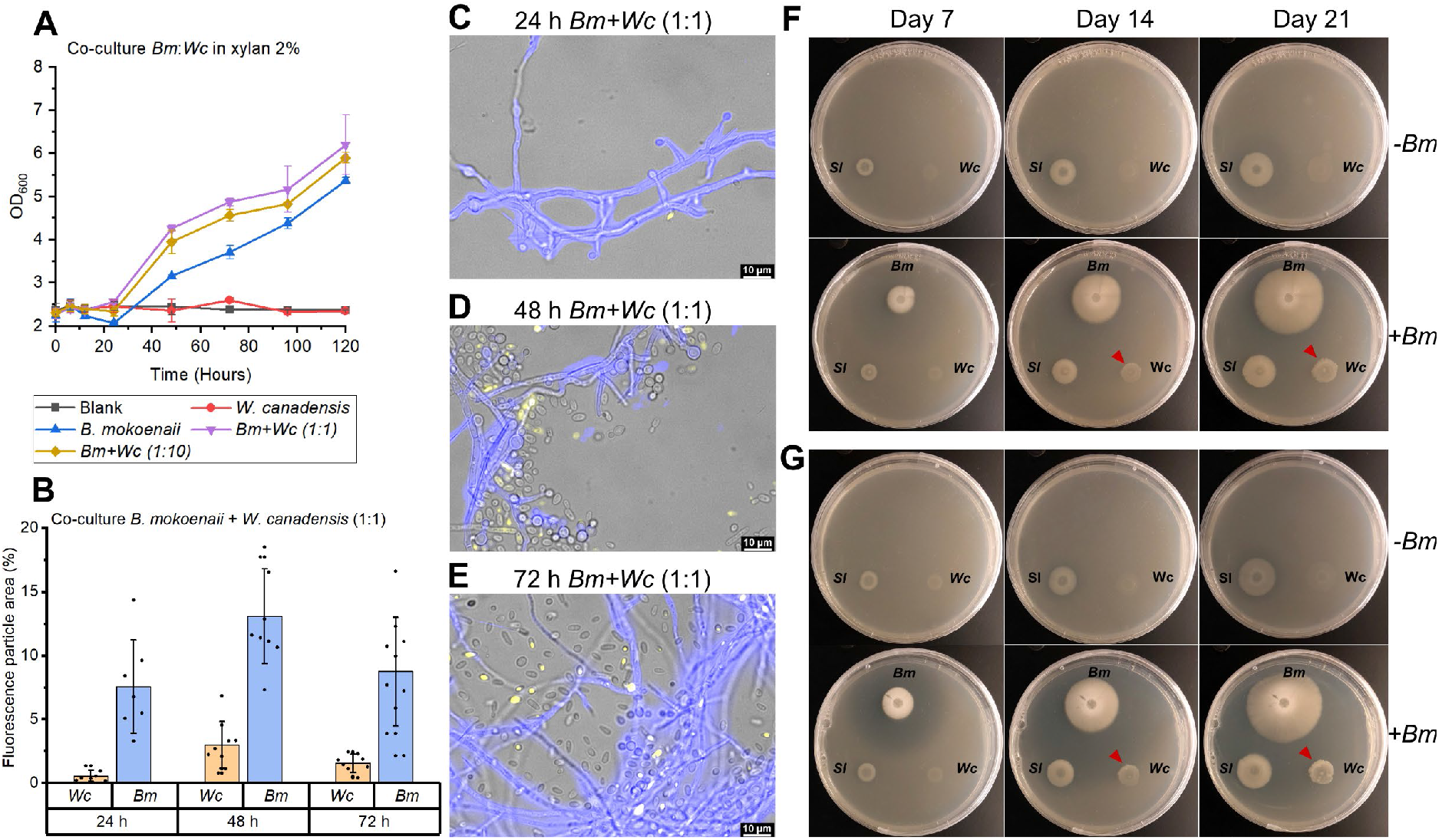
Co-cultures of yeast growth in xylan. (A) Co-cultures of *B. mokoenaii* and *W. canadensis* at different starting ratios in 2 % beechwood glucuronoxylan in triplicates. (B) Distribution in fluorescence particle area (%) of co-cultures of *B. mokoenaii* (blue) and *W. canadensis* (yellow) at an initial ratio of 1:1. (C) Representative images from fluorescence microscopy of *B. mokoenaii* and *W. canadensis* (1:1 ratio) after 24 h, (D) after 48 h, and (E) after 72 h. Co-cultures on agar plates with 0.4 % beechwood glucuronoxylan and (F) or 0.4 % wheat arabinoxylan (G) over time with or without the addition of *B. mokoenaii* (*Bm* +/-). The clearing zones correlate with xylanase-mediated xylooligosaccharide release and enables *W. canadensis* growth (red arrows). *Bm* = *Blastobotrys mokoenaii*, *Sl* = *Scheffersomyces lignosus*, *Wc*=*Wickerhamomyces canadensis*.

### Biochemical characterization of putative *endo*-1,4-β-xylanases

As all three species showed capability to metabolize xylan, we sought to characterize the likely responsible enzymes. The selection of putative xylanases was based on CAZyme profile predictions, secretome in-gel proteomics from xylan cultures (10), and preliminary flow cell proteomics targeting cell-associated proteins in beechwood GX cultures (unpublished results). The candidates were from the major xylanase-containing families GH11 (*Bm*Xyn11A), GH30 (*Bm*GH30_7) in *B. mokoenaii*, and GH10 (*Sl*Xyn10A) in *S. lignosus*. As mentioned, *W. canadensis* lacks obvious xylanase candidates but encodes enzymes from GH5, a large polyspecific family with many different β-1,4-cleaving enzymes (including xylanases), which are further grouped into subfamilies (3, 13). *W. canadensis* encodes enzymes from subfamily 9 (GH5_9), 12, 22, and 49. Of these, only GH5_22 has been shown to contain xylanolytic enzymes, though no reported activity of GH5_49 members exist. Thus, the GH5_22 (*Wc*Xyn5_22A) and GH5_49 (*Wc*Xyn5_49A) enzymes were selected as potential xylanases from this species. Codon optimized genes were synthesized and all targets except *Bm*GH30_7 were successfully expressed in *Pichia pastoris* and purified (gene sequences in Supplemental Information, List S1 and purified recombinant enzymes in Fig. S2). Each enzyme was assayed at appropriate dilutions (Fig. S3), temperatures (40-55 °C) and pH (5–6) (Fig. S4) corresponding to the growth preferences of each species.

The specific activities of the purified enzymes were determined on various polysaccharides, biomasses, and synthetic substrates; *Bm*Xyn11A, *Sl*Xyn10A, *Wc*Xyn5_49A, and *Wc*Xyn5_22A all showed *endo*-β-1,4-xylanase activity on beechwood GX and wheat AX but no or minor activity on other carbohydrates (Table 1). *Bm*Xyn11A had the highest specific activity of the recombinant enzymes on xylan (4350 U·mg^-1^), nearly fourfold higher on beechwood GX compared to the closely related GH11 XynB from *Aspergillus niger An76* (1146 U·mg^−1^) (14). *Sl*Xyn10A was active on both beechwood GX and wheat AX, though with relatively low activity (13-15 U mg^-1^) compared to many bacterial and fungal xylanases (15). *Wc*Xyn5_49A similarly had moderate activity (Table 1), though no direct comparisons within the subfamily can be made as it to the best of our knowledge is the first member biochemically characterized. While *Wc*Xyn5_49A was mainly active on xylan, *Wc*Xyn5_22A showed activity on both *p*NP-β-D-xyloside and xylans, albeit low, indicating both β-xylosidase activity and *endo*-β-1,4-xylanase activity. β-xylosidase activity was previously reported for *Pc*GH5_22 from the fungus *Phanerochaete chrysosporium* (16). Although we cannot rule out that the studied yeasts possess additional *endo*-xylanases, likely including *Bm*GH30_7 and a second GH10 copy in *S. lignosus* and possibly other yet undiscovered enzymes, these results clearly demonstrate that *Bm*Xyn11A, *Sl*Xyn10A and *Wc*Xyn5_22A and *Wc*Xyn5_49A all are active on GX and AX.

**Table 1.**
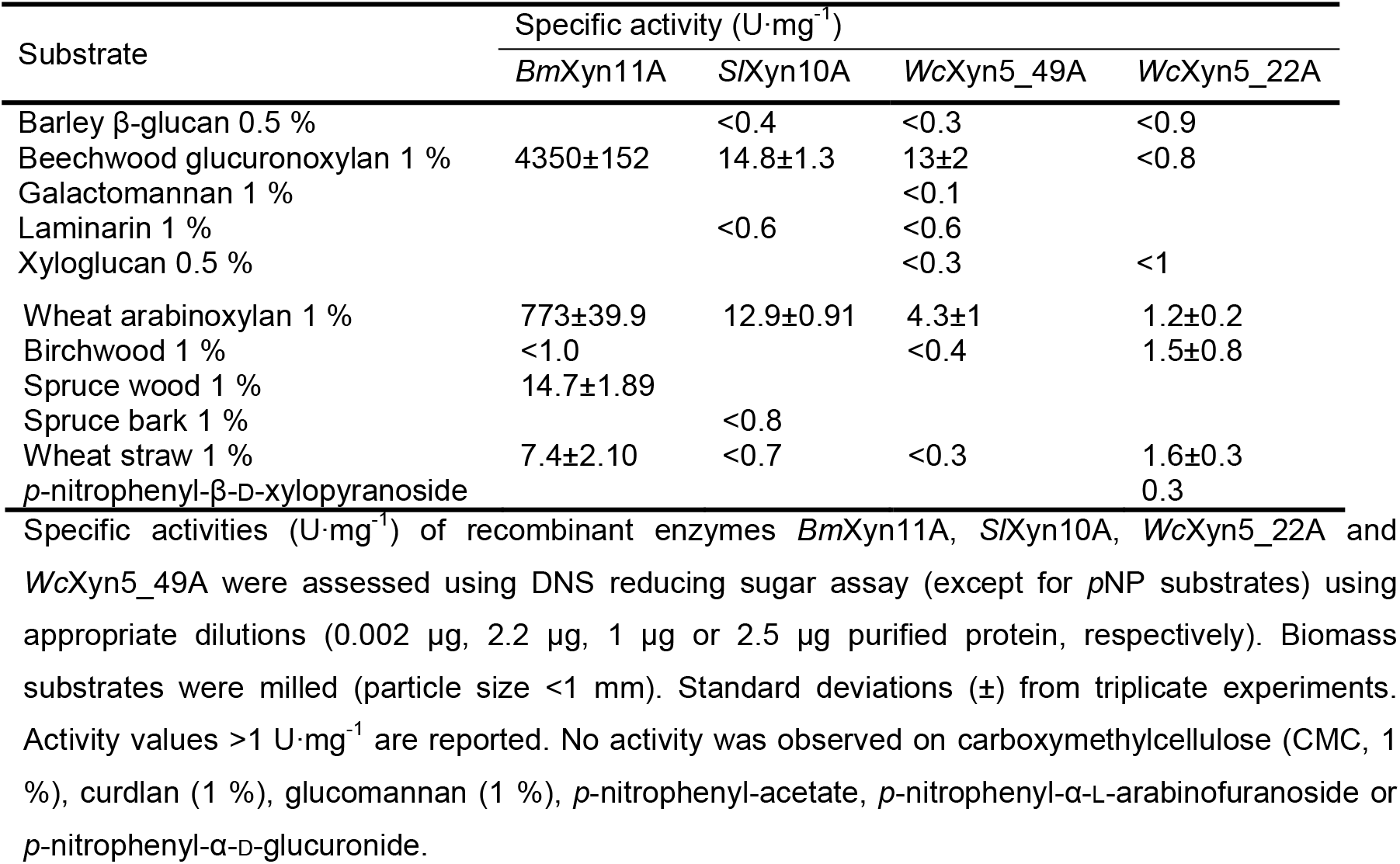
Specific activity of recombinant yeast enzymes.

### Xylan deconstruction and xylooligosaccharide profiles of recombinant xylanases

To compare and further characterize the identified yeast xylanases, degradation of beechwood GX and wheat AX and their hydrolysis products formed over time were determined using an enzyme concentration of 0.1 μM. After 24 h, XO profiles were analyzed by HPAEC-PAD (Fig. 4A-B). *Bm*Xyn11A reached near-maximum xylan hydrolysis levels after only 15 min (Fig. 4A), releasing mainly smaller XOs (xylose, X1; xylobiose, X2; and xylotriose, X3) (Fig. 4C). S*l*Xyn10A took longer (8 h) to reach near-maximum hydrolysis levels of both xylans and yielded mainly X2 and X3 from AX, and also longer XOs from GX (similar levels of X2-xylohexaose, X6). The differences in XO distributions between *Bm*Xyn11A and *Sl*Xyn10A from GX corresponds to previous studies, where GH11 xylanases require three unsubstituted xylose units for attack on the xylan backbone while GH10 xylanases can accommodate more branched xylans (17, 18). *Wc*Xyn5_49A generated mainly X2, X3 and xylotetraose (X4) from GX but also xylopentaose (X5) from wheat AX, while *Wc*Xyn5_22A generated minor amounts of XOs, indicating that it is likely not the main xylanase of *W. canadensis* (Fig. 4B).

**Figure 4.**
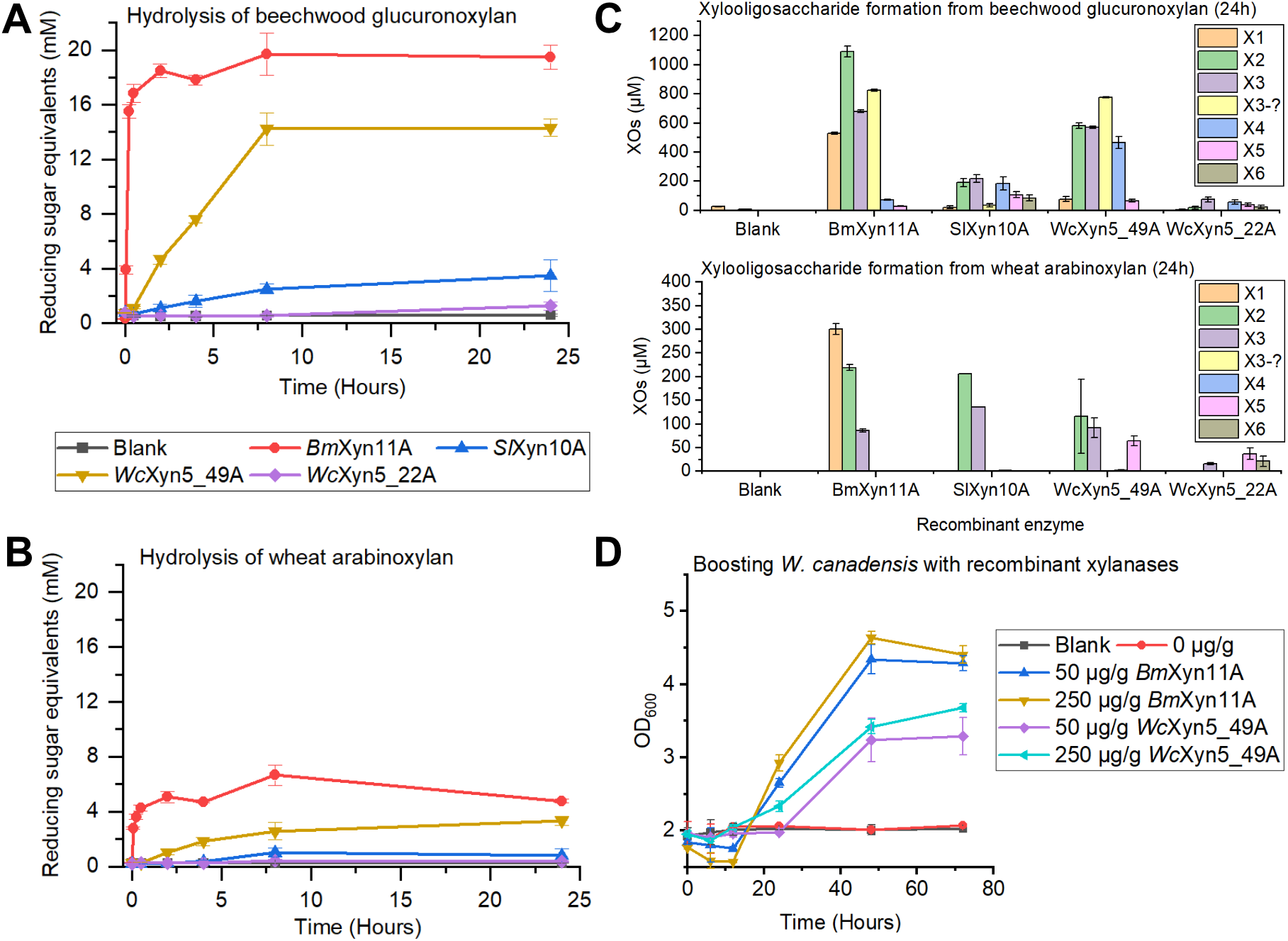
Effect of xylanase treatment on beechwood glucuronoxylan and wheat arabinoxylan and boosting of yeast growth. (A) Hydrolysis of beechwood glucuronoxylan and (B) wheat arabinoxylan by recombinant enzymes over 24 h using 0.1 μM enzyme in triplicates. (C) Xylooligosaccharide formation from beechwood glucuronoxylan and wheat arabinoxylan after 24 h in duplicates. Xylose (X1), xylobiose (X2), xylotriose (X3), xylotetraose (X4), xylopentaose (X5), xylohexaose (X6). XOs= Xylooligosaccharide. (D) Boosting of *W. canadensis* growth in 2 % beechwood glucuronoxylan by supplying the cultures with *Bm*Xyn11A and *Wc*Xyn5_49A, in lower (50 μg/g xylan) and higher (250 μg/g xylan) concentrations in biological triplicates.

To provide further experimental support for the endo-xylanase activity of *Wc*Xyn5_49A, we supplemented *W. canadensis* xylan cultures with different amounts of recombinant *Wc*Xyn5_49 as well as *Bm*Xyn11A (positive control). Addition of either of the recombinant xylanases resulted in an initial drop followed by a significant increase in OD_600_, indicative of xylan solubilization and cellular growth, respectively (Fig. 4D). Although addition of *Bm*Xyn11A resulted in higher growth rates and final OD_600_ compared to *Wc*Xyn5_49A, the results firmly establish *Wc*Xyn5_49A as an endo-xylanase that can successfully generate XOs and trigger *W. canadensis* growth in xylan.

### Xylanase structural analyses

We aimed to solve the structures of the recombinantly expressed xylanases for comparisons with previously solved xylanase structures. We successfully crystallized and solved the structure of *Bm*Xyn11A extending to 1.55 Å resolution, by molecular replacement using an AlphaFold2-predicted structure of *Bm*Xyn11A as search model. Data have been deposited in the PDB with accession code 8B8E, and collection and refinement statistics are shown in Table S1. The structure of the *P. pastoris-expressed Bm*Xyn11A is the first GH11 structure from a budding yeast. The overall structure adopted the expected GH11 β-jelly roll fold (Fig. 5A), and the asymmetric unit contained five protein molecules (Fig. S5). By superposition of *Bm*Xyn11A with the structure of an *Aspergillus niger* GH11 xylanase in complex with X5 (PDB code 2QZ2 (19), the *Bm*Xyn11A catalytic residues were identified as Glu113 (nucleophile) and Glu204 (proton donor) (Fig. 5B and C). Comparisons with previously solved GH11 members from bacteria and fungi shows high conservation, even among kingdoms (Fig. S6), with Cα-RMSD values < 0.6 Å to *Bm*Xyn11A, which could suggest either retention of this enzyme in *B. mokoenaii* after the split between yeast and filamentous fungi or possibly a recent horizontal gene transfer event.

**Figure 5.**
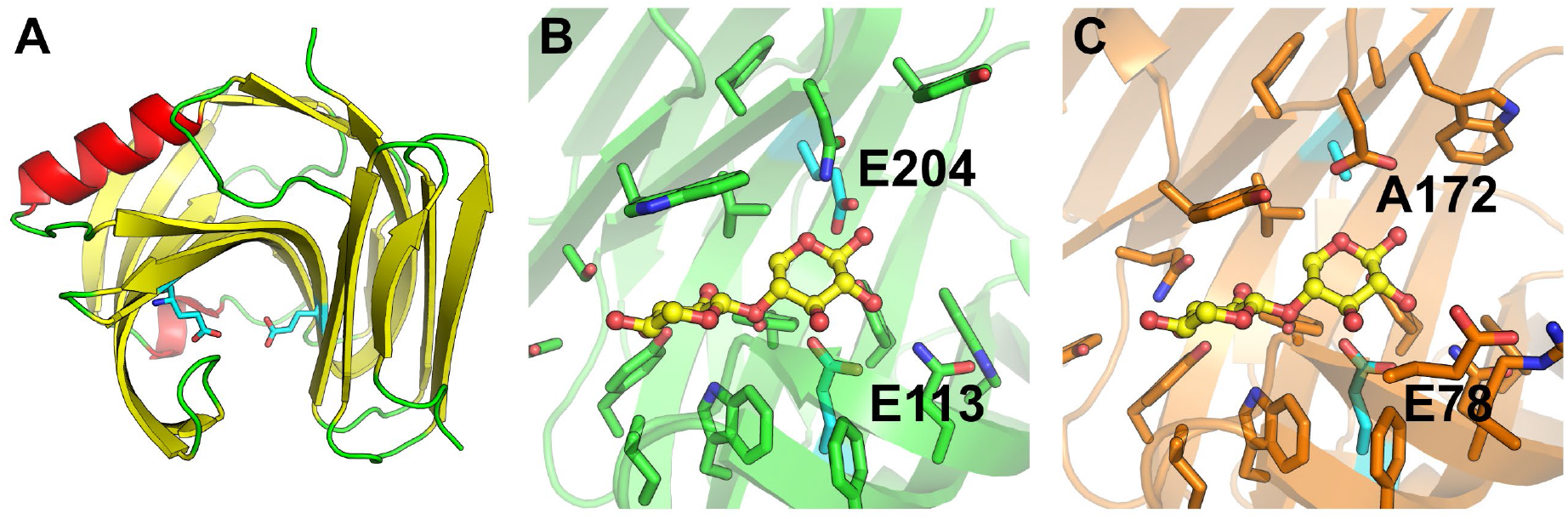
Crystal structure of *Bm*Xyn11A and comparison with other GH11 members. (A) The overall fold of *Bm*Xyn11A (chain A), with β-sheets in yellow, α-helices in red, loops in green, and catalytic Glu residues as cyan sticks. (B) Closeup of the active site cleft of *Bm*Xyn11A (green) superimposed with the bound oligosaccharide in yellow ball and stick representation derived from the *Aspergillus niger* GH11 xylanase (PDB: 2QZ2, inactivated EA variant, crystallized with X5), shown in the same orientation in (C).

While we were unable to experimentally solve the structures of the other yeast xylanases, we could predict their structures using AlphaFold2 (20) (Fig. 6). *Sl*Xyn10A conformed to other GH10 enzymes, with Cα-RMSD value of 0.937 to *Cb*Xyn10C from *Caldicellulosiruptor bescii* (PDB 5OFK), which enabled identification of the catalytic residues of *Sl*Xyn10A: E163 (proton donor) and E270 (nucleophile) (21). The active site appears to be an extended cleft, which suggest an ability to accommodate long xylan chains or XOs, and likely substitutions depending on the angle of the backbone xylose moieties (Fig. 6A). The active sites of *Sl*Xyn10A and *Cb*Xyn10C are nearly identical in each sugar-binding subsite, and infers six or more subsites in *Sl*Xyn10A (21). *Wc*Xyn5_22A and *Wc*Xyn5_49A both adopted the expected GH5 (α/β)_8_ barrel, though *Wc*Xyn5_49A in addition was appended with a C-terminal bundle of four alpha helices (Fig. 6, top left, green; residues 400-end) of unknown function; it did not show any similarity to known carbohydrate binding module (CBM) families (3, 22). Such helical bundle inserts have been shown to confer thermal stability in other proteins (23, 24), but does not appear to be the case for *Wc*Xyn5_49A as it did not exhibit significant thermal stability (Fig. S4). By comparison of the GH5 enzymes to XEG5A (PDB: 4W89 (25)) the catalytic residues could be identified for *Wc*Xyn5_49A as Glu223 (proton donor) and Glu327 (nucleophile) (Fig. 6B); and for *Wc*Xyn5_22A as Glu191 (proton donor) and Glu294 (nucleophile) (Fig. 6C). The active sites of *Sl*Xyn10A and *Wc*Xyn5_49A adopted expected clefts able to accommodate extended polysaccharide substrates (Fig. 6), but a helical insert outside the active site of *Wc*Xyn5_22A creates a large ridge that appears to block positive subsites (towards the reducing end). Such apparent blockage of the active site could suggest *exo*-activity rather than *endo*, though this does not correlate with the biochemical analyses (Fig. 4), and the modelled positioning of this insert may not be of biological relevance despite high confidence scores (Fig. S7). In contrast, *Wc*Xyn5_49A possesses a highly accessible active-site cleft, suggesting it can accommodate large substrates (Fig. 6). Future experimental determination or modelling of ligand-bound structures of the yeast xylanases may reveal deeper molecular information about their specificities.

**Figure 6.**
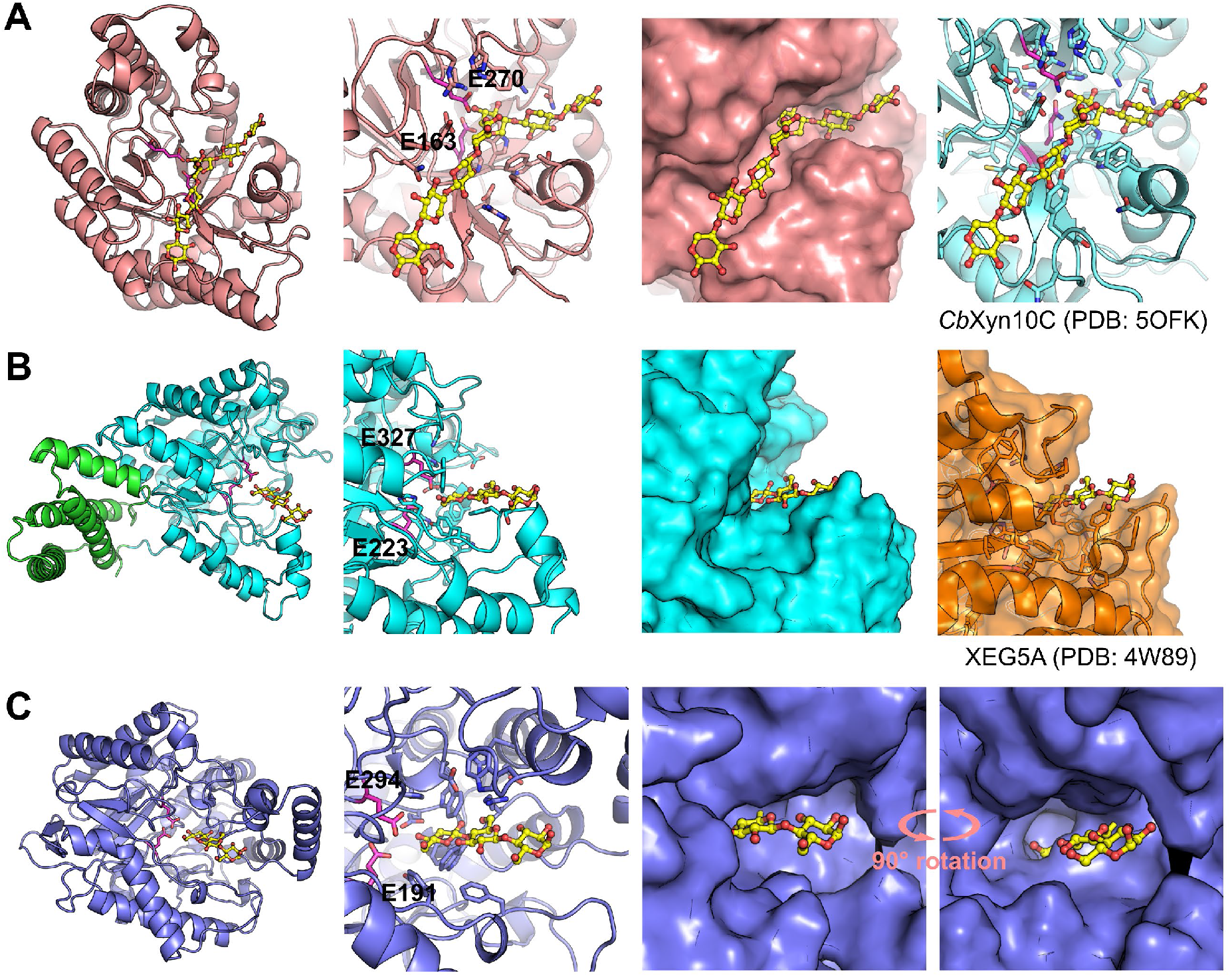
Predicted AlphaFold2 structures for GH5 and GH10 xylanases and β-xylosidases characterized in this study. (A) *Sl*Xyn10A. (B) *Wc*Xyn5_49A. (C) *Wc*Xyn5_22A. Left: Overview of each enzyme. The helical bundle in *Wc*Xyn5_49A is shown in green sticks. The catalytic Glu residues are magenta sticks. In the GH5 structures, a superimposed oligo from the structure of XEG5A (PDB: 4W89) is shown in yellow sticks; in the GH10 structure the oligo is superimposed from *Cb*Xyn10C (PDB: 5OFK). Second from left: closeups of each active site. Second from right: surface representation, highlighting the different shapes of the binding clefts. In (C), a 90° rotated view reveals the deep cleft of *Wc*Xyn5_22A with limited surface accessibility. Figure S7. visualizes pLDDT scores for each predicted model.

## Discussion

Our previous work using comparative genomics among known xylan-degrading budding yeasts revealed different setups of xylanolytic enzymes, where many species lack xylanases from typical GH families (10). The CAZyme family GH5 has over 50 subfamilies that mainly contain enzymes cleaving various β-(1,4)-linked oligo- and polysaccharides (3), and therefore, stood out as a likely source of endo-β-1,4-xylanases in the yeasts without obvious candidates (2). In this study, we demonstrate experimentally that three xylanolytic yeasts, *B. mokoenaii*, *S. lignosus* and *W. canadensis*, isolated from different environments and belonging to different phylogenetic clades, have evolved distinct strategies to degrade xylan that manifests as different growth behaviors on this carbon source. We also provide experimental validation of xylanase activity for the yeast enzymes *Bm*Xyn11A, *Sl*Xyn10A, *Wc*Xyn5_22A and *Wc*Xyn5_49A, the latter being from a previously uncharacterized CAZyme subfamily.

Interestingly, GH5_49 appears enriched in yeast compared to filamentous fungi and bacteria (3). In the dataset of 332 genome sequenced budding yeasts, 319 have predicted GH5_49 genes and 126 possess GH5_22 genes (10), for which there has previously been no (GH5_49) or little (GH5_22) experimental characterization. Although 16 of the 24 known xylanolytic yeasts in the dataset contain both GH5_22 and GH5_49 genes, there are also many yeasts that have such subfamily genes but still are not known to depolymerize and grow on xylan (3). It is clear that additional characterization of GH5_22 and GH5_49 members is needed to verify whether they generally act as xylanases or have different functions in other yeast and non-yeast species. Exploring possible synergies with other CAZymes would further shed light on their biological role(s).

In the three species investigated here, also other enzymes may have roles in xylan turnover, but our physiological and biochemical analyses begin to paint a picture of their highly divergent xylanolytic strategies (Fig. 7). *B. mokoenaii* is the standout species, much thanks to its secreted powerful *Bm*Xyn11A (10) which likely facilitates the rapid growth on xylan. Also, the pseudo-mycelial growth morphology of *B. mokoenaii* may be an advantage in its native soil environment, similar to many filamentous fungi (26). Yeasts that possess GH11 xylanases seem rare, and we have only been able to find one other ascomycetous yeast, *Aureobasidium pullulans*, and a few basidiomycetous yeasts containing GH11 genes, all of which display superior capacity to degrade xylan (2, 27). In contrast to *B. mokoenaii* and *W. canadensis*, the xylanase activity of *S. lignosus* was mainly localized to the cell surface, which is also a feature of the β-xylosidase activity of all three species. Possibly, the native environment of *S. lignosus* in the presumably highly competitive insect gut has exerted a selective pressure for more selfish nutrient acquisition behavior, where full secretion of xylanases into the extracellular environment does not promote fitness. Similar ‘selfish’ behavior of microorganisms has been demonstrated for example in the human gut, where certain polysaccharide-metabolizing bacteria tether both enzymes and carbohydrate-binding proteins to the cell surface to limit leakage of smaller sugars (28, 29). Intracellular enzymes require transporters that can internalize the substrate first, which are uncommon for larger oligosaccharides (30). The surface-located β-xylosidase activity of the three yeasts studied here would however suggest that mainly small sugars are transported over the plasma membranes.

**Figure 7.**
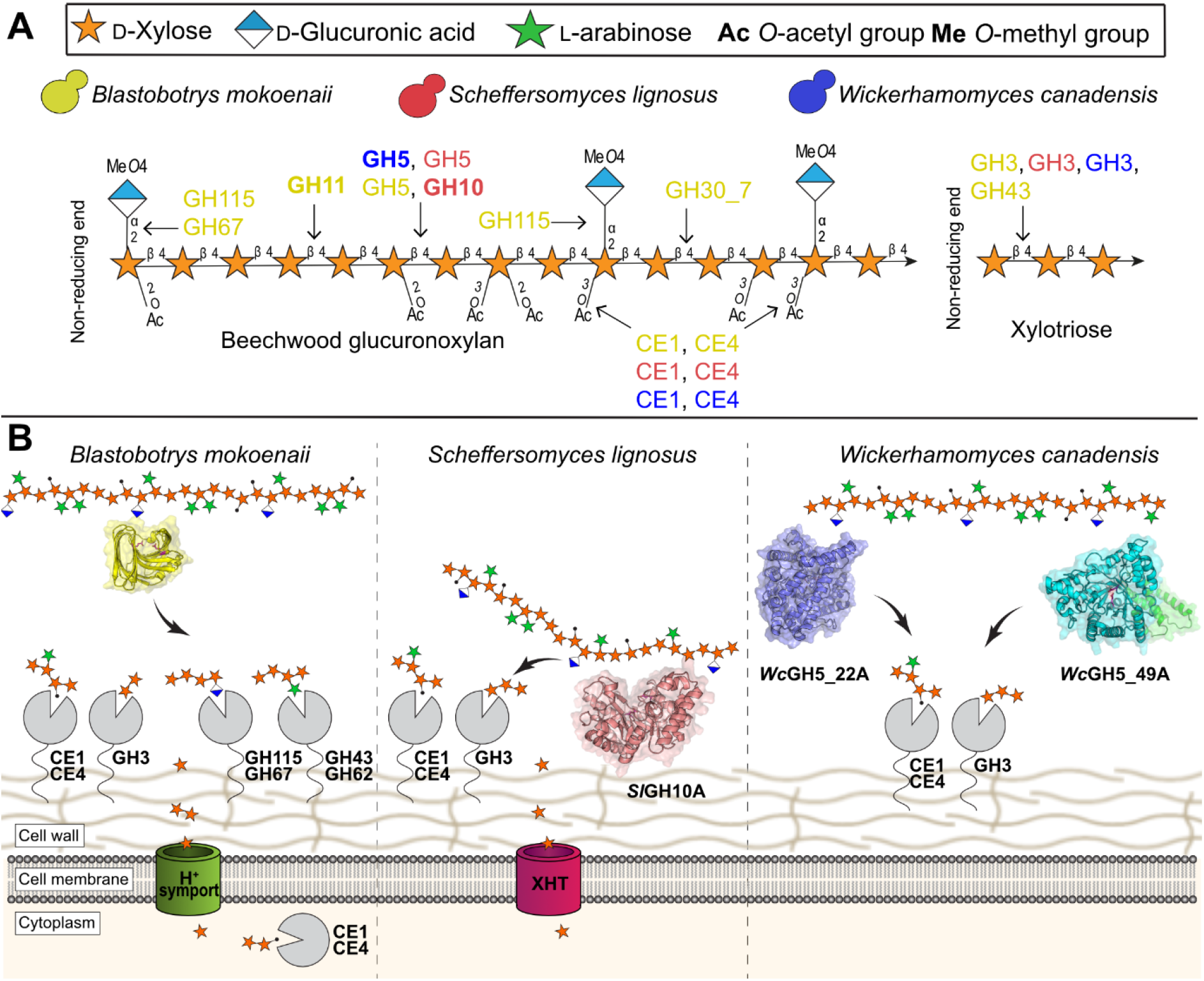
Schematic view of the xylanolytic strategies of the three investigated species. (A) Predicted xylanolytic CAZymes (10) of *Blastobotrys mokoenaii, Scheffersomyces lignosus* and *Wickerhamomyces canadensis* represented by color (yellow, red and blue, respectively). Arrows from individual enzyme families indicate enzyme activity in beechwood glucuronoxylan degradation. CE= carbohydrate esterase; GH= glycoside hydrolase. (B) Xylan, *B. mokoenaii* utilizes a secreted xylanase which depolymerizes xylan away from the cells and liberates oligosaccharides in the process, *S. lignosus* and *W. canadensis* instead maintains the xylanase activity close to the cell surface, which might give a competitive advantage in their respective environment. Putative enzymes are shown in gray with CAZy family memberships indicated. Acetyl groups are shown as black dots. XHT = xylose/glucose transporters.

Interestingly, we observed that *W. canadensis*, even when pre-grown on xylose, cannot grow on the beechwood GX used in this study. Apparently, it lacks the ability to sense intact xylan polymers, but its xylanolytic machinery was inducible by small amounts of XOs, or by addition of xylanases that presumably lead to the same products. This phenomenon has also been observed previously for some other xylanolytic yeasts (31). It is possible that *W. canadensis* can sense longer oligosaccharides than tested here, similar to some bacteria which require medium- to large-sized xylan fragments to induce *endo*-β-1,4-xylanase expression (32). In its natural habitat, it is thus likely that *W. canadensis* needs neighboring primary degraders of xylan releasing XOs for it to kickstart its own xylanolytic machinery. Biomass-degrading microbes are not presumed to act alone in nature, but in concert with other species, by either collaborating or competing for the sugars. In this sense, *W. canadensis* seems to have an opportunistic xylanolytic strategy, which possibly could be advantageous by not expending resources for constitutive xylanase production and instead responding only when xylan is being degraded nearby. It is interesting to note that *W. canadensis* displays relatively high β-xylosidase activities, which may suggest a competitive advantage for XOs released during active xylan degradation and a successful xylan scavenging behavior.

Our findings on xylanolytic yeasts may provide a starting point for further fundamental research on their biomass-degrading enzymes and cellular physiology. Future characterization of other species will further reveal how commonplace the proposed biological role(s) of the here studied yeast are in global biomass decay ecosystems. Co-culturing species from the same, or similar, environments would be of high interest to compare with our *B. mokoenaii* – *W. canadensis* cultures, as such experiments could showcase whether other opportunistic species like *W. canadensis* can exhibit similar behavior also on other types of polysaccharides. The findings in this study are also of industrial importance. Mono- and co-cultures of xylanolytic yeasts can be used in consolidated bioprocesses, where simultaneous hydrolysis of biomass/xylan and conversion into product are carried out by the microorganisms rather than the former being performed by expensive enzyme cocktails (33). The yeasts characterized here might be used as future cell factories in their own right, or as donors of genes to already established yeast cell factories based on e.g. *Saccharomyces cerevisiae*. Moreover, the enzymes studied here may also find application in food and feed production, or the pulp and paper industries. For example, *Bm*Xyn11A showed no activity on CMC which is a biotechnologically attractive trait in pulp bleaching (34).

In summary, our study shows that between dissimilar yeast species, there are wide-ranging degradation strategies and enzymes. These may be more successfully transferred to a yeast host such as *S. cerevisiae* than the enzymes from filamentous fungi and bacteria used today. We conclude that knowledge of new xylanolytic enzymes and strategies may help scientists to find new strategies for metabolic engineering, cell-factory design, and synthetic biology of yeasts in the future.

## Materials & Methods

### Yeast growth characterization in xylan

#### Liquid cultures

Yeasts strains were ordered from the ARS Culture Collection, USA (NRRL; https://nrrl.ncaur.usda.gov/). Yeast growth in beechwood glucuronoxylan or xylose was determined by inoculating yeasts with a starting OD_600_= 0.1 in 10 mL Delft minimal media (5 g L^−1^ ammonium sulfate, 3 g L^−1^ potassium phosphate, 1 g L^−1^ magnesium sulfate, vitamins and trace metals as described previously (35) additionally containing either 20 g L^−1^ (w/v) of beechwood glucuronoxylan (Megazyme, Ireland) or 20 g L^−1^ xylose (w/v). Cultures were kept at either 30 °C (for *B. mokoenaii* CBS 8435, Y-27120) or room temperature (~21 °C) for *S. lignosus* (CBS 4705, Y-12856) and *W. canadensis* (CBS 1992, Y-1888) at 150 rpm for 96 h in 100 mL baffled flasks. Xylooligosaccharides (<95% XOs) from corncob (Carlroth, Germany) were added at 2 g L^−1^ (w/v) to kickstart *W. canadensis* growth, with or without additional xylan. Yeasts were pre-cultured in Delft minimal media with 20 g L^−1^ xylose and washed in Delft minimal media without carbon source before inoculation into xylan or xylose. For co-cultures, yeast precultures (Delft minimal media with 20 g L^−1^ xylose) were inoculated separately with a starting OD_600_= 0.1 or in different ratios (1:1 and 1:10) in Delft medium + 2% beechwood glucuronoxylan (Megazyme, Ireland) for 120 h at RT, 200 rpm. For growth boosting with recombinant enzymes, 50 μg/g xylan or 250 μg/g xylan enzymes was supplemented to cultures.

### Yeast subcellular enzyme activities

To quantify the subcellular *endo*-β-1,4-xylanase activity during xylan degradation (secretome, cell-associated or intracellular), cells were harvested by centrifugation (3000×g, 15 min) and divided into two fractions. Each cell fraction was washed in Delft minimal medium (without carbon source), one fraction was kept for cell-associated activity, and the other (OD_600_= 5) was lysed by eight cycles of bead beating at 8000 rpm, 30 s, followed by addition of Y-PER (Yeast Protein Extraction Reagent; Pierce, Rockford, IL, USA) breaking buffer with 25 mM 1,4-dithiothreitol. Lysis efficiency was monitored microscopically. The soluble intracellular fraction was isolated by centrifugation (13,000×g, 5 min) and assayed alongside the cell-free supernatant (secretome), and the intact cell pellets, respectively. The assay contained 10 g L^−1^ wheat AX or beechwood GX and 50 mM sodium acetate buffer (pH 5.5) added to cell pellets or 25 μL cell-free supernatants mixed in a 96-well plate. The mixture was incubated at 30 °C for 30 min followed by immediate chilling on ice. Reducing sugar ends were determined by the dinitrosalicylic acid (DNS) method (Miller, 1959). All enzymatic measurements were performed in triplicates. For de-branching activities, cell fractions were incubated with either 2.5 mM *p*-nitrophenyl-acetate, *p*-nitrophenyl-α-L-arabinofuranoside, *p*-nitrophenyl-α-D-glucuronide or *p*-nitrophenyl-β-D-xylopyranoside in 200 μL reactions containing 20 mM sodium phosphate (pH 7). The reactions were incubated at 30 °C for 20 min at 350 rpm in 96-well plates, which were then centrifuged at 4 °C (4000×g, 5 min) to remove intact yeast cells, and 100 μL was transferred to a new 96-well plate, where *p*-nitrophenol was quantified at 405 nm over 30 min.

### Bright field and Fluorescent microscopy of yeast co-cultures

Yeast cells from xylan mono- and co-cultures were sampled (100 μL) at different timepoints (24, 48, 72 or 96 h), centrifuged (10.000×g, 5 min), washed in 1 mL 10 mM HEPES buffer pH 7.2. Co-cultures were stained with 0.15 μL FUN-1 and 0.5 μL Calcoflour white in 100 μL 10 mM HEPES buffer pH 7.2 + 2% glucose using a LIVE/DEAD™ Yeast Viability Kit (ThermoFisher, Germany) for 30 min in dark at RT. Fluorescence images was recorded using a Leica DFC360 FX microscope (Germany) with a 100x-oil-immersion objective and a DFC 360 FX camera. Excitation filters for YFP (480-520 nm) was used for FUN-1 using an exposure time of 75 ms with 1.8 gain and excitation filter A4 (320-400 nm) was used for Calcoflour white using an exposure time of 5 ms and 1.8 gain. Emission spectrums were designated yellow and blue colors for FUN-1 and Calcoflour white, respectively, in the LAS X (Leica) software. Total fluorescent particle area and pixel intensity values in the 12-bit images (1392×1040 pixels) were used to differentiate and quantify blue and yellow fluorescence using the Image J (Fiji) software.

### Agar plate xylan sensing assay

10 μL washed yeast pre-cultures (Delft minimal media with 20 g L^−1^ xylose) with a starting OD_600_= 5 were pipetted onto Delft minimal medium agar plates (2 %) containing 0.4 % beechwood glucuronoxylan (Megazyme, Ireland) or wheat arabinoxylan (Megazyme, Ireland) with appropriate distance between strains. Plates were incubated at room temperature for 21 days and pictures were taken daily to follow xylan clearing zones and yeast colony growth.

### Cloning and protein production

Genes encoding the putative proteins *Bm*Xyn11A, *Bm*GH30_7, *Sl*Xyn10A, *Sl*GH5_22, *Wc*Xyn5_22A and *Wc*Xyn5_49A (sequence IDs in Supplemental Information) were codon optimized and synthesized by Twist Bioscience (USA) for heterologous expression in *P. pastoris* X-33 and cloned into pPICZα A vectors using EasySelect (Thermofisher, Germany). Native signal peptides were removed from *Bm*Xyn11A and *SlXyn10A*. The vector was digested with *EcoRI* and *Sall* and inserts ligated by T4 ligase. Recombinant pPICZα A constructs contained yeast α-secretion factor, candidate gene and His6-tag and transformed into *E. coli* DH5α One Shot Top10 cells (Invitrogen). Transformants were selected based on Zeocin resistance (25 μg mL^−1^) confirmed by colony PCR. Vector propagation was performed in 1.5 mL LB (low salt + Zeocin 75 μg mL^−1^) and GeneJET PCR Purification Kit (ThermoFisher, Germany). *P. pastoris* cells were transformed using ~50 ug linearized pPICZα A by electroporation and clones were selected using Zeocin (150 μg mL^−1^) on YPD plates with 1 M sorbitol while colony PCR confirmed presence of recombinant vectors. *P. pastoris* clones were grown in small scale (5 mL) to confirm recombinant enzyme activity before scale-up to 1L baffled flasks at 28 °C in 400 mL rich buffered glycerol-complex medium (BMGY). Methanol (1%, v/v) was used to induce expression in buffered methanol-complex medium (BMMY) as described in the EasySelect Pichia Expression kit, and the cells grown for six days as previously described (37).

### Enzyme purification

Proteins were purified by immobilized metal affinity chromatography (IMAC) using Ni-Sepharose excel resin (GE Healthcare, USA). The column was first washed with 5 column volumes of loading buffer (50 mM tris(hydroxymethyl)aminomethane (TRIS), pH 8, 250 mM NaCl). After loading and washing of bound proteins, they were eluted using the same buffer with additional 250 mM imidazole. *Sl*Xyn10A was de-glycosylated with endo H (NEB, USA) according to the supplier’s protocol. *Wc*Xyn5_22A and *Wc*Xyn5_49A were further purified using anion exchange chromatography on a HiTrap SP HP column (Cytiva) using 50 mM TRIS (pH 8) as loading buffer and elution using a linear gradient with 0-1 M NaCl. Protein purity was evaluated by SDS-PAGE. A Nanodrop 2000 spectrophotometer (Thermo Fisher Scientific, Germany) was used to determine protein concentration using the predicted values for molecular weights and extinction coefficients (Expasy ProtParam server).

### Biochemical characterization

*Endo*-β-1,4-xylanase activity was assayed using a 200 μL mixture of 10 g L^−1^ beechwood GX (Megazyme, Ireland) in 50 mM sodium acetate buffer (pH 5.5) using 10 μL purified enzyme at suitable dilutions (Fig. S3). The mixture was incubated for 30 min followed by immediate chilling on ice and inactivation at 98 °C for 5 min. For time-course measurements, 50-100 μL aliquots were sampled at 0, 0.5, 2, 4, 8, and 24 h and immediately heat-inactivated and stored at −20 °C, before determining reducing sugar ends using the DNS method. For pH and temperature optimum measurements, the following buffers were used, at 100 mM: sodium acetate pH 5 or pH 5.5, sodium phosphate pH 6-8, sodium citrate pH 3-4, 2-(N-morpholino)ethanesulfonic acid (MES) pH 5-6 and *N*-cyclohexyl-2-aminoethanesulfonic acid (CHES) pH 8-10. In other polysaccharides including: wheat arabinoxylan (Megazyme, Ireland), birchwood glucuronoxylan (Sigma-Aldrich, Germany), xyloglucan (tamarind, Megazyme, Ireland), mixed-linkage β-1,3/1,4-glucan (barley, Megazyme, Ireland), galactomannan (guar/locust bean gum, Sigma-Aldrich, Germany), glucomannan (konjac, Sigma-Aldrich, Germany), curdlan (Merck, USA), pectin (citrus, Sigma-Aldrich, Germany), laminarim (*Laminaria digitata*, Sigma-Aldrich, Germany), carboxymethyl cellulose (Sigma-Aldrich, Germany) and the ball-milled (~40 μm) local Swedish biomasses: spruce wood, birchwood, wheat straw and spruce bark, reducing sugar ends released by purified recombinant enzymes was determined by the DNS method. Specific activity on *p*NP-acetate, *p*NP-β-xyloside, *p*NP-α-L-arabinofuranoside and *p*NP-α-glucuronide (2.5 mM) was measured using 0.002 μg (*Bm*Xyn11A), 2.2 μg (*Sl*Xyn10A), 1 μg (*Wc*Xyn5_22A) or 2.5 μg (*Wc*Xyn5_49A) purified protein, was determined in 20 min reactions as described earlier.

### Xylooligosaccharide analysis by ion chromatography

XO formation from hydrolysis of beechwood glucuronoxylan (Megazyme, Ireland) and wheat arabinoxylan (Megazyme, Ireland) by recombinant enzymes was analyzed after 24 h reactions (0.1 μM enzyme, using 1 % xylan, at 40 °C in 1000 μL total volume). 100 μL aliquots were inactivated by boiling, diluted in water, centrifuged (5 min, 13,000 rpm) and the supernatants were filtered (0.22 μm) before storage at 4 °C until analysis by high performance anion-exchange chromatography coupled with pulsed amperometric detection (HPAEC-PAD) using an ICS-5000 system (Dionex, USA) at 25 °C. Separation of hydrolysis products was performed with a CarboPac PA200 (250 mm × 3 mm) column (Thermofisher, Germany) and the following eluents A: Water: B: 300 mM sodium hydroxide and C: 100 mM sodium hydroxide and 1 M sodium acetate using a flow rate of 0.5 mL min^−1^. The mobile phase gradient can be found in Table S2. Standards (5-1000 μM) of xylose (X1), xylobiose (X2), xylotriose (X3), xylotetraose (X4) xylopentaose (X5) and xylohexaose (X6) (Megazyme, Ireland) were used for quantitation.

### Protein crystallography, structure determination and molecular modelling

Prior to crystallization, a sample of *Bm*Xyn11A was de-glycosylated using Endo H (RT, overnight). This sample as well as the *P. pastoris*-expressed untreated enzyme were further purified using size-exclusion chromatography, using a HiLoad 16/600 Superdex 200 pg size exclusion column (Cytiva), operated by an ÄKTA Explorer FPLC using 25 mM Tris pH 8.0 as buffer. Comparison with a calibration curve of protein standards confirmed the enzyme was monomeric in both cases and showed a small degree of glycosylation in the case of the untreated enzyme. The enzymes were concentrated using Amicon 10 kDa cutoff centrifugal filter units to 65 mg mL^−1^ for the untreated *Bm*Xyn11A, and 12 mg mL^−1^ for the Endo H treated. Both proteins were subjected to crystallization screening using a Mosquito robot in sitting-drop vapor diffusion trays at 20 °C, with drops of 0.3 μL protein + 0.3 μL reservoir solution, 40 mL reservoir volume, using the JCSG+, Morpheus, and PACT Premier screening kits (Molecular Dimensions). Crystals were not obtained for the Endo H-treated enzyme. For the untreated *Bm*Xyn11A, rod-shaped crystals were obtained in 2 weeks from condition D12 from JCSG+ (0.04 M potassium phosphate monobasic, 16 % w/v PEG 8000, 20% w/v glycerol; 65 mg/mL *Bm*Xyn11A). Optimized, large crystals were obtained in 3 weeks from the same condition with 32.5 mg/mL *Bm*Xyn11A. These were flash-frozen in liquid nitrogen. Diffraction data were obtained on 22^nd^ April 2022 on the BioMAX beamline at MAX IV (Lund, Sweden) at 100 K. The dataset was collected at 0.1° increments over 360°, processed using Mosflm (38), and the structure solved by molecular replacement with Phaser in the Phenix program suite (39), using a AlphaFold2 model of *Bm*Xyn11A as the search model (20, 40). The solvent content indicated two molecules in the asymmetric unit. The protein structure was manually rebuilt after phasing using Coot (41), and refined using Phenix Refine. R_free_ was monitored using 5 % of the diffraction data selected at random prior to refinement. Detailed data collection and refinement statistics are outlined in Table S1. The coordinates for the structure were validated and deposited in the PDB with accession code 8B8E. Structure prediction models of *Wc*Xyn5_22A and *Wc*Xyn5_49A were produced using the AlphaFold2 software (20, 40).

## Acknowledgements

The authors would like to thank the ARS Culture Collection for providing yeast cultures. We acknowledge MAX IV Laboratory for time on Beamline BioMAX under Proposal 20200093. Research conducted at MAX IV, a Swedish national user facility, is supported by the Swedish Research council under contract 2018-07152, the Swedish Governmental Agency for Innovation Systems under contract 2018-04969, and Formas under contract 2019-02496. The AlphaFold2 structure predictions were enabled by resources provided by the Swedish National Infrastructure for Computing (SNIC) at Chalmers Centre for Computational Science and Engineering, partially funded by the Swedish Research Council through grant agreement no. 2018-05973.

## Authors’ contributions

CG, JL and JR conceived the project, CG, JL and JR designed the experiments; JR, ARS and TC performed the experiments. JR performed growth cultures, enzymatic assays, recombinant protein expression. ARS and JR performed protein purification and characterization. TC performed crystallography and AlphaFold predictions together with JL. JR, ARS and TC interpreted the data and wrote the manuscript. JR, CG, JL, ARS and TC revised the manuscript. All authors read and approved the final manuscript.

## Declarations

Ethics approval and consent to participate – Not applicable.

## Consent for publication

All authors consented on the publication of this work.

